# Neuronal progenitors suffer genotoxic stress in the *Drosophila* clock mutant *per^0^*

**DOI:** 10.1101/2024.08.13.607601

**Authors:** Nunzia Colonna Romano, Marcella Marchetti, Anna Marangoni, Laura Leo, Diletta Retrosi, Ezio Rosato, Laura Fanti

## Abstract

The physiological role and the molecular architecture of the circadian clock in fully developed organisms are well established. Yet, we have a limited understanding about the function of the clock during ontogenesis. We have used a null mutant (*per^0^*) of the clock gene *period* (*per*) in *Drosophila melanogaster* to ask whether PER may play a role during normal brain development. In 3^rd^ instar larvae, we have observed that absence of functional *per* results in increased genotoxic stress compared to wild type controls. We have detected increased double strand DNA breaks in the central nervous system and chromosome aberrations in dividing neuronal precursor cells. We have demonstrated that reactive oxygen species (ROS) are causal to the genotoxic effect and that expression of PER in glia is necessary and sufficient to suppress such a phenotype. Finally, preliminary evidence indicates that absence of PER may result in less condensed chromatin, which may contribute to DNA damage.

## Introduction

The circadian clock is a timing mechanism that tunes biochemistry and physiology to the environment. In *Drosophila melanogaster* flies, the clock revolves around the expression of two genes, *period* (*per*) and *timeless* (*tim*), that are regulated by their own protein products. Rhythmic expression starts during the day with the transcription of *per* and *tim* by CLOCK/CYCLE (CLK/CYC), a heterodimeric transcription factor. After synthesis, PER and TIM undergo a maturation process that begins in the early evening comprising dimerization, progressive post-translational modifications, and accumulation. Late at night, the two proteins become nuclear and competent inhibitors of CLK/CYC. However, PER and TIM modifications trigger their degradation, which releases the inhibition on CLK/CYC. Thus, during the day, *per* and *tim* transcription starts again, beginning a new cycle^1^. Mammals have a circadian clock also, which is fundamentally similar to the one in *Drosophila*^2^. Importantly, CLK/CYC and the homologous CLK/BMAL1 in mammals, directly and indirectly, control the expression of a large part of the genome^3-7^. This suggests that the clock and/or its constituents may be important regulators of chromatin structure^8-12^.

There is evidence that components of the circadian clock are expressed during early ontogenesis, but the role that they, or the clock as a process, play in development is unclear^13-16^. There is a well-known interdependence between circadian clock and metabolism in fully developed organisms^17^. Interestingly, metabolic reprogramming is both cause and effect of changes in differentiation status during development^18^. This suggests that the circadian clock, or its parts, may be involved in the developmental programme of the organism.

In this study we start exploring the role PER may have in the development of the nervous system in *Drosophila*. Mutants *per*-null (*per^0^*) grow to adulthood, are fertile and do not show gross morphological abnormalities. Nevertheless, *per^0^* flies have neurological defects as shown by alterations in circadian rhythms, memory formation and sleep architecture^19^. Additionally, they exhibit mild anatomical defects such as an irregular location of a group of neuroendocrine cells in the brain and an abnormal arborization pattern of a cluster of clock neurons^20-22^. Overall, these observations suggest that PER may be required during development for the correct assembly of neuronal circuits. Furthermore, as *per^0^* flies have metabolic defects such as reduced mitochondrial function, increased sensitivity to reactive oxygen species (ROS) and shorter lifespan, there may be a link between metabolic and neurological dysfunctions^23-27^.

Our working hypothesis is that lack of PER, *via* abnormal metabolism, may cause genotoxic stress impacting the developmental programme of the nervous system. Third instar larvae constitute an informative model since the presence of hundreds of mitotic cells allows to measure, in addition to DNA damage, chromosomal aberrations in dividing neuronal precursors.

In flies, the formation of the central nervous system (CNS) proceeds through two ontogenetic phases. During early embryogenesis neuronal stem cells called neuroblasts (NBs) delaminate from the embryonic neuroectoderm, become larger and starts dividing. The transition between embryonic and larval development results in the NBs becoming quiescent (with some exceptions). The first larval stage marks the beginning of feeding. The availability of nutrients triggers the re-activation of the cell-cycle in NBs. In each brain lobe (BL) there are approximatively 100 type I NBs (NBs I). At each division, they generate a NB I and a ganglion mother cell (GMC) that divides once to produce neurons and/or glia. NBs II are much fewer, 8 per BL, but undergo a remarkable amplification of their lineages. At every division a NB II generates another NB II and an intermediate nuclear progenitor cell (INP). The INP is initially immature, but after maturation it divides 5-6 times, each producing one proliferating INP and one GMC that divides once to give rise to neurons/glia^28, 29^. The VNC contains only NBs I. Thus, in the CNS of 3^rd^ instar larvae there are hundreds of mitotic cells at any one time, which is ideal for assessing chromosome integrity in neuronal precursors (Fig. S1 linked to Fig. 1).

**Fig 1.**
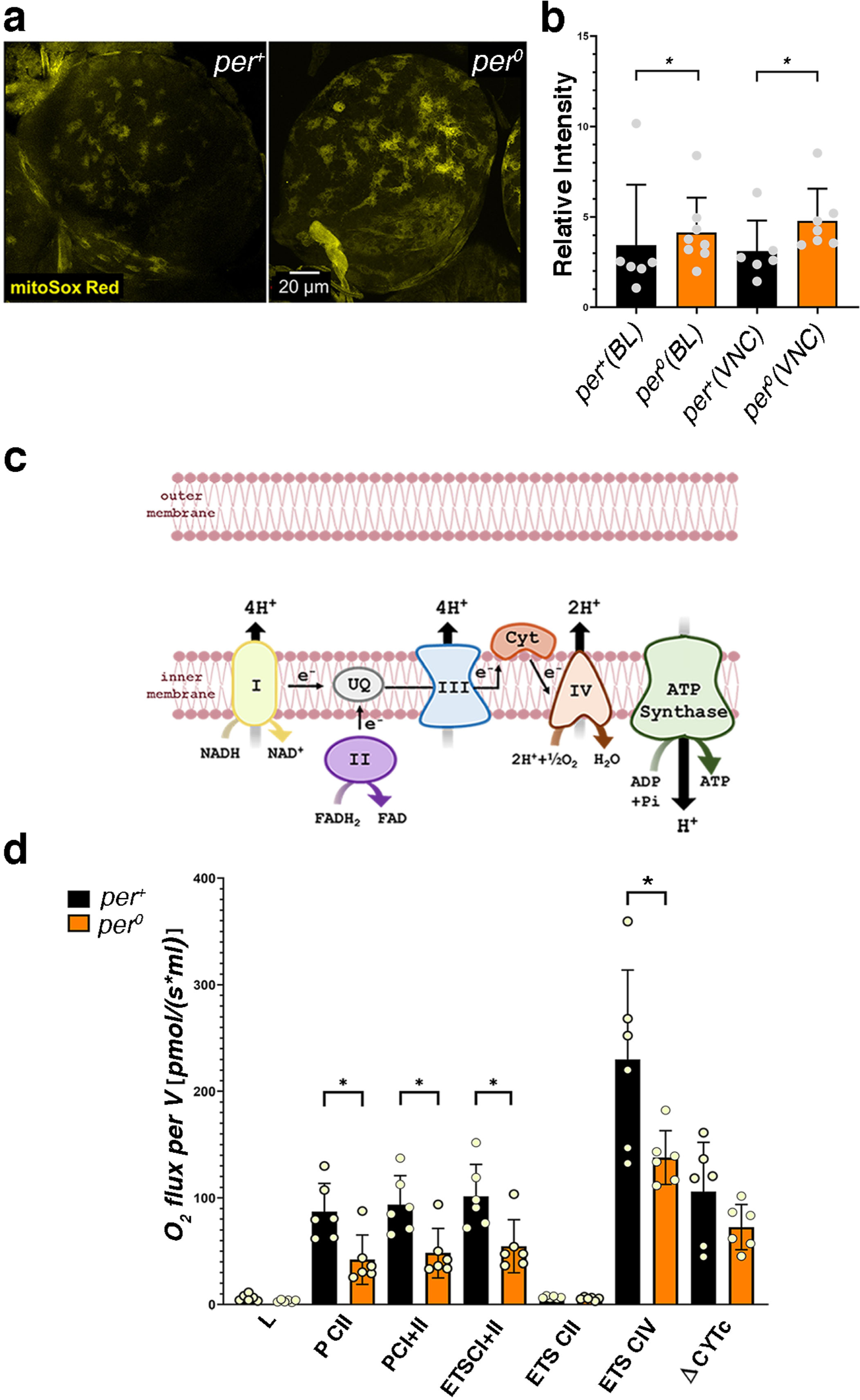
High levels of ROS and mitochondrial respiration defects in *per^0^* mutants. a) Brain lobes stained with mitoSox Red in *per*^+^ and *per^0^* 3^rd^ instar larvae. Confocal maximum intensity projections. Size bar = 20 μm. ZT = 2. b) Quantification of mitoSox Red signal in *per*^+^ and *per^0^*. Brain lobes (Kolmogorov-Smirnov test, *P=0.042) and ventral cords (Kolmogorov-Smirnov test, *P=0.015) were compared separately. Points show individual samples. Error bars = SD (standard deviation). ZT=2. c) Cartoon of the mitochondrial complexes involved in oxidative phosphorylation (OXPHOS). d) High resolution respirometry (oxygen flux per volume [pmol·s^-1^·mL^-1^]) in 3-5 day old *per*^+^ and *per^0^* flies. OXPHOS capacity related to complex I (P CI; Mann-Whitney, *P=0.026) and complex I plus II (P CI + II; Mann-Whitney, *P=0.026) were reduced in *per^0^* compared to *per*^+^ controls. Additionally, *per^0^* showed reduced electron transfer capacity through complex I plus II (ETS CI + II; Mann-Whitney, *P=0.026), and through complex IV (ETS CIV; Mann-Whitney, *P=0.041) but not complex II alone (ETS CII; Mann-Whitney, P=0.132). There were no differences in mitochondrial leak (L; Mann-Whitney, P=0.093) and in mitochondrial integrity (ΔCYTc; Mann-Whitney, P=0.310) between the two genotypes. Points show individual samples. Error bars = SD. ZT=1. All samples were males obtained by reciprocal crossing (♀ *CS* x ♂ *per*^0^ and *vice versa*).

In this report, we use third instar larvae to show that lack of PER results in DNA damage and in a high frequency of chromosome aberrations in dividing neuronal precursor cells. These genotoxic effects correlate with a rise in ROS levels. Additionally, we provide evidence that PER expression in glia is necessary and sufficient to avoid chromosome aberrations. Finally, we establish a link between PER-dependent genotoxic phenotypes and defects in chromatin architecture. We suggest that PER, either on its own or as part of the clock, controls chromatin states by regulating the metabolic programme of the cells and that such a regulation is important for normal development.

## Results

### per^0^ MUTANTS ARE SUBJECT TO A HIGH ROS BURDEN

Previous reports have shown that *per^0^* flies have abnormal metabolism and are sensitive to ROS^25^. We used MitoSOX Red, a fluorogenic superoxide indicator dye that is targeted to the mitochondria, to measure ROS levels in the CNS of larvae (at ZT2, *Zeitgeber* Time 2, corresponding to 2h after lights on). We detected higher levels of fluorescence in *per^0^* compared to *per*^+^, indicating that the mutant is subject to a higher ROS burden than the control (Fig. 1a, b). We used high resolution respirometry (Oroboros oxygraph) on whole larva extracts (at ZT1) to identify defects in the mitochondrial complexes and the electron transport chain (Fig. 1c). Surprisingly, we did not uncover any difference in respiration between *per^0^* and *per*^+^ larvae (Fig. S2 linked to Fig. 1). We do not have a clear explanation for this finding. Perhaps, lower and/or uncoupled respiration (the dissipation of the mitochondrial H^+^ gradient), which are regular features of larval mitochondria, mask differences between the wild type and the mutant^30, 31^. Thus, we measured respiration in adult fly extracts (at ZT1). We detected a reduction in the oxidative phosphorylation (OXPHOS) capacity related to complex I and complex I plus II in *per^0^*compared to *per*^+^ controls. Additionally, per^0^ showed reduced electron transfer capacity through complex I plus II and through complex IV (Fig. 1d). A decrease in electron transfer capacity may cause over-reduction of the ubiquinone pool and formation of superoxide radicals. In summary, we have identified significant respiration defects in *per^0^* flies that may explain the observed ROS increase in larvae.

### THE CNS OF per^0^ LARVAE SHOWS DNA DAMAGE AND CHROMOSOME ABERRATIONS

Cells exposed to high levels of ROS are prone to oxidative damage, which includes double strand DNA (dsDNA) breaks^32^. Phosphorylation of histone variant H2AV at serine 137 (referred to as γ-H2AV) marks the recognition of dsDNA breaks and the promotion of repair^33^. Thus, we asked whether *per^0^* larvae may show higher anti-γ-H2AV immune reactivity than *Canton-S* (*CS*) wild type controls. We produced CNS-squash preparations (at ZT 2) and we labelled them with anti-γ-H2AV (Fig. 2a). As expected, the number of immune positive cells was higher in *per^0^* than in *CS* (Fig2. b). Notably, we obtained similar results using whole mount preparations and larvae that had a more homogenous genetic background. For the latter, we exploited the fact that the *per* locus is X-linked; thus, we selected the male progeny of reciprocal crosses [♀ *per*^0^ x ♂ *CS* and *vice versa*]. These results are illustrated in the following section.

**Fig 2.**
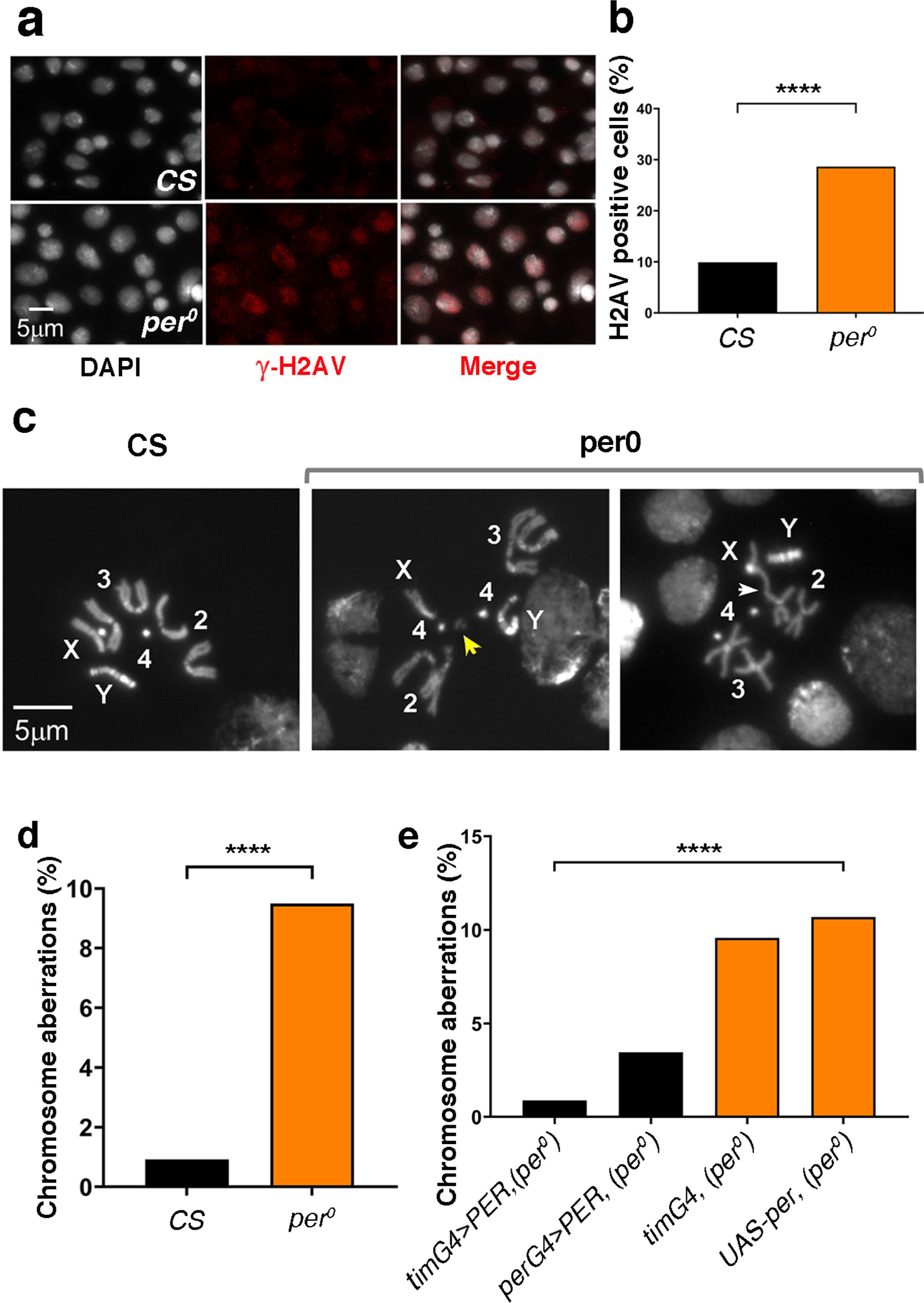
DNA damage and chromosome aberrations in *per^0^* mutants. a) Anti-γ-H2AV immune labelling (red) in CNS-squash preparations from 3^rd^ instar male larvae. DNA is stained with DAPI (white). Size bar = 5 μm. ZT=1. b) Proportion of cells showing anti-γ-H2AV immune signal in CNS-squash preparations from *CS* and *per^0^*3^rd^ instar male larvae. Fisher’s exact test, ****P<0.0001. Total number of cells, N = 1323, 985, respectively. ZT=1. c) Mitotic metaphases in CNS-squash preparations from 3^rd^ instar larvae. Left, *CS*, showing a normal metaphase. Right, chromosome aberration in *per^0^*. The yellow arrow indicates a chromosome fragment (break). The white arrow points to a fusion. 2, 3, 4 identify the autosomes. X, Y label the sex chromosomes. Size bar = 5 μm. ZT=2. d) Frequency of chromosome aberrations in *CS* and *per^0^*. Fisher’s exact test, ****P<0.0001. ZT=2. Total number of metaphases scored, N = 436, 527, respectively. e) PER overexpression rescues chromosome aberrations in *per^0^*. The overexpression of PER (*UASper16*) using the pan-circadian *per-* and *tim-GAL4* drivers drastically reduced the frequency of aberrations in an otherwise *per^0^* background. Chi-square= 49.50, df=3, ****P<0.0001. Total number of metaphases scored (from left to right), N = 337, 463, 345, 1021. ZT=2.

dsDNA breaks may lead to chromosome aberrations (breaks, fusions, or translocations, see Fig. 2c). We calculated the proportion of mitotic metaphases showing chromosome aberrations in another set of CNS-squash preparations (at ZT2). Indeed, we observed a significantly higher proportion of aberrations in *per^0^* than *CS* larvae (Fig. 2d). Then, we tested (at ZT2) whether overexpressing PER (*UAS-per16*) in *per^0^* larvae, using *perGAL4* or *timGAL4* as a driver, could rescue the high rate of chromosome defects. Larvae *per^0^*expressing either *perGAL4*>PER or *timGAL4*>PER showed wild type levels of chromosome aberrations (Fig. 2e). Since *per* and *tim* define circadian cells when co-expressed, this finding may suggest that it is a defective clock, rather than the specific lack of PER, being causal to the genotoxic phenotype.

### BUFFERING OF ROS RESCUES DNA DAMAGE AND CHROMOSOME ABERRATIONS IN per^0^

Vitamin C is a ROS-scavenger without negative side-effects in *Drosophila*^34^. Larvae *per^0^* that had developed on medium supplemented with vitamin C (40mM) showed a significant reduction in dsDNA breaks (anti-γ-H2AV signal) both in whole mount and in CNS-squash preparations (Fig. 3a-d). Chromosome aberrations were likewise reduced (Fig. 3e). The expression of *Ciona intestinalis* alternative oxidase (AOX) lessens the production of mitochondrial ROS. AOX avoids the overload of the electron transport chain by accepting electrons directly from the ubiquinone pool and reducing O_2_ into H_2_O^35^ (Fig. 3f). In *per^0^*larvae we expressed AOX in all putative clock cells (*timGAL4*>AOX), which resulted in rescue of chromosome aberrations (Fig. 3g).

**Fig 3.**
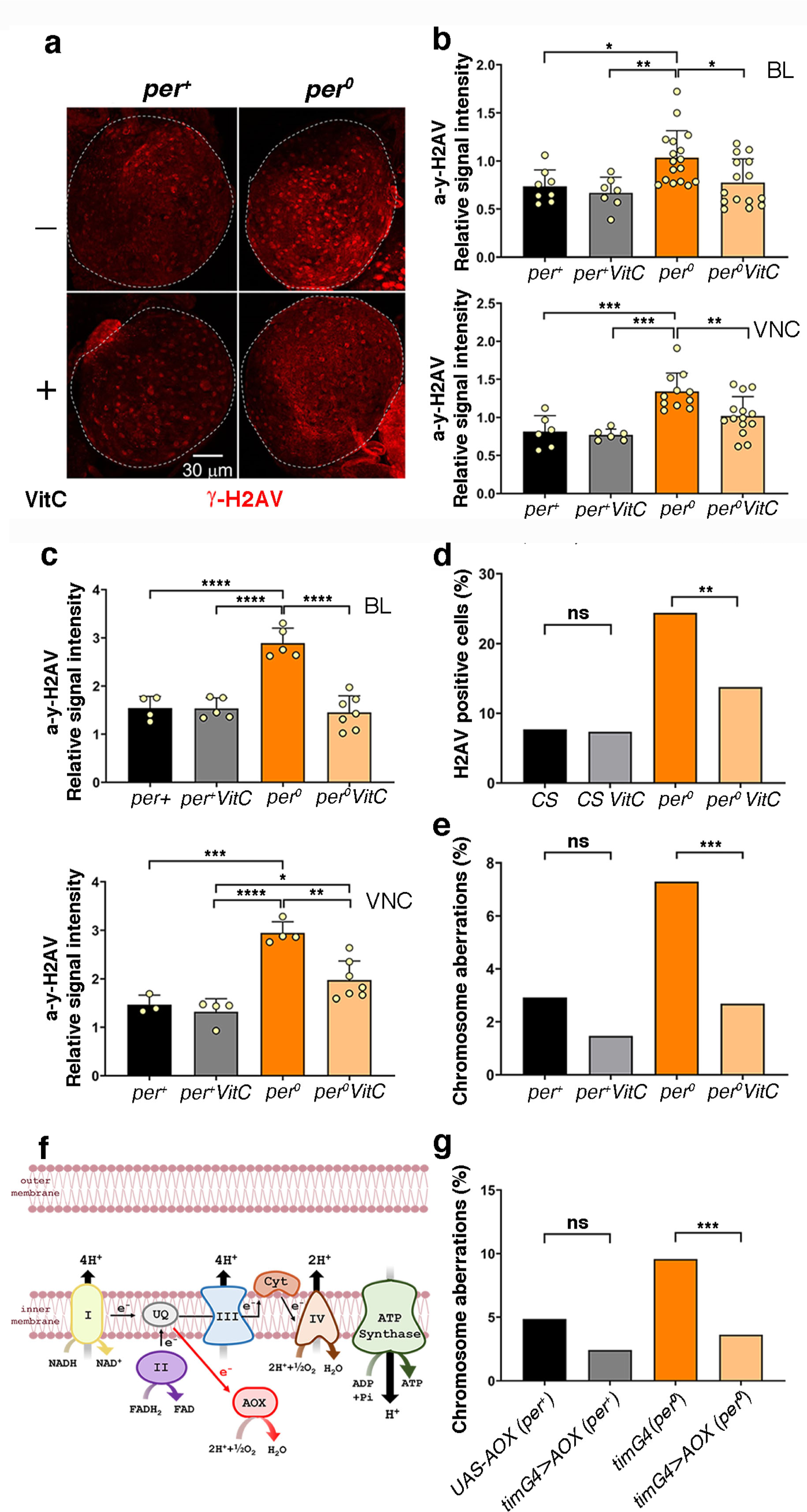
ROS buffering and reduction decrease DNA damage and chromosome aberrations in *per^0^*. Treatment with vitamin C (VitC, 40 mM in standard medium, from embryo) reduced anti-γ-H2AV immune labelling (a-d) and chromosome aberrations (e) in *per^0^*3^rd^ instar larva males (a-e). a) Brain lobes (BLs) from 3^rd^ instar *per*^+^ and *per^0^* male larvae obtained by reciprocal crossing (♀ *CS* x ♂ *per*^0^ and *vice versa*); anti-γ-H2AV on whole mount. Confocal maximum intensity projections. Dashed lines outline BLs. Size bar = 30 μm. ZT=2. b, c) Quantification of anti-γ-H2AV immune fluorescence intensity [relative signal intensity = (signal-background)/background] in whole mount CNS from 3^rd^ instar *per*^+^ and *per^0^* male larvae as above. b & c show two independent experiments (imaged on different microscopes). Normal distribution of data was confirmed with the Shapiro-Wilk test. b) Top, BL (one per individual). Two-way ANOVA, Genotype (F_1, 43_=7.318, **P=0.0097), VitC treatment (F_1, 43_=4.732, *P=0.0352), Genotype x VitC treatment (F_1, 43_=1.617, P=0.2104). Tukey’s multiple comparisons test, *per*^+^ *vs*. *per^0^*, *P=0.0282; *per*^+^ VitC *vs*. *per^0^*, **P=0.0075; *per*^0^ *vs*. *per^0^* VitC, *P=0.0196. Bottom, VNC. Two-way ANOVA, Genotype (F_1, 33_=23.75, ****P<0.0001), VitC treatment (F_1, 33_=5.241, *P=0.0286), Genotype x VitC treatment (F_1, 33_=3.044, P=0.0903). Tukey’s multiple comparisons test, *per*^+^ *vs*. *per^0^*, ***P=0.0003; *per*^+^ VitC *vs*. *per^0^*, ***P=0.0001; *per*^0^ *vs*. *per^0^* VitC, **P=0.0066. ZT=1. c) Top, BL (one per individual). Two-way ANOVA, Genotype (F_1, 14_=21.03, ***P=0.0004), VitC treatment (F_1, 14_=40.05, ****P<0.0001), Genotype x VitC treatment (F_1, 14_=39.53, ****P<0.0001). Tukey’s multiple comparisons test, *per*^+^ *vs*. *per^0^*, ****P<0.0001; *per*^+^ VitC *vs*. *per^0^*, ****P<0.0001; *per*^0^ *vs*. *per^0^* VitC, ****P<0.0001. Bottom, VNC. Two-way ANOVA, Genotype (F_1, 14_=46.94, ****P<0.0001), VitC treatment (F_1, 14_=12.97, **P=0.0029), Genotype x VitC treatment (F_1, 14_=7.040, *P=0.0189). Tukey’s multiple comparisons test, *per*^+^ *vs*. *per^0^*, ***P=0.0001; *per*^+^ VitC *vs*. *per^0^*, ****P<0.0001; *per*^+^ VitC *vs*. *per^0^*, *P=0.0236; *per*^0^ *vs*. *per^0^* VitC, **P=0.0011. Points show individual samples. Error bars = SD. ZT=1. d) Proportion of γ-H2AV immune positive cells in CNS squash preparations from 3^rd^ instar male larvae. Treatment with VitC did not affect *CS* (Fisher’s exact test, P=0.9209) but lowered the proportion of γ-H2AV immune labelled cells in *per^0^* (Fisher’s exact test, **P=0.0011). Total number of cells scored (from left to right), N = 912, 775, 295, 298. ZT=2. e) Proportion of chromosome aberrations in CNS squash preparations from 3^rd^ instar *per*^+^ and *per^0^*male larvae obtained by reciprocal crossing (♀ *CS* x ♂ *per*^0^ and *vice versa*). Treatment with VitC lowered the proportion of chromosome aberrations. The effect was small in *per*^+^ (Fisher’s exact test, P=0.1886) but highly significant in *per^0^*(Fisher’s exact test, ***P<0.0005). Total number of metaphases scored (from left to right), N = 468, 440, 439, 633. ZT=1. AOX overexpression rescues chromosome aberrations (f, g). f) Cartoon showing the position of the Alternative Oxidase (AOX) in the electron transport chain. g) The overexpression of AOX using the pan-circadian *tim-GAL4* driver reduced the frequency of aberrations. The effect was marginal in *per*^+^ (Fisher’s exact test, P=0.0663) but highly significant in *per^0^*(Fisher’s exact test, ***P<0.0009). Total number of metaphases scored (from left to right), N = 397, 387, 456, 412. ZT=1. Males obtained by reciprocal crossing [♀ *UAS-AOX (per^+^)* x ♂ *tim-GAL4* (*per*^0^) and *vice versa*].

### PER IS EXPRESSED IN GLIA, INCLUDING CORTEX

In third instar larvae there are 9 neurons per brain lobe (BL) showing robust and rhythmic PER and TIM expression. These are *bona fide* clock neurons and correspond to clusters recognised in the adult brain. We asked whether PER may be present, additionally, in diving cells such as NBs and GMCs. The transcription factor PROSPERO (PROS) is asymmetrically distributed in the cytoplasm of NBs I, whereas it becomes nuclear in GMCs^28^. We carried out α-PER and α-PROS immune staining at ZT23, a time when PER is generally nuclear, but failed to identify double-labelling in either type of dividing cells (Fig. 4a, b). Instead, we observed a weak and diffuse α-PER immune signal around α-PROS immune reactive cells (Fig. 4a, b). The NBs and their progeny are enveloped by cortex glia. These are large, web-like cells, born during the mid-embryonic stage, that surround and support the NBs and their lineages functioning as ‘niche’^36^. Thus, we carried out α-PER and α-REVERSE POLARITY (α-REPO) immune staining (at ZT23); the latter labels the nuclei of all glia cells. Comparing *per^0^* and *per*^+^ larvae further suggested that PER may be expressed, albeit weakly, in the cytoplasm of cortex glia cells (Fig. 4c, d). To verify that such a weak α-PER immune reactivity identifies cortex glia, we took advantage of a ‘classic’ PER reporter that accumulates in cells. It consists of a genomic fragment of *per*, including the promoter region and up to the first half of the encoded protein, that is fused in frame to the bacterial *lacZ* gene. The fusion protein, SG, is stable and does not cycle^37^. We employed *repo-GAL4* to express GFP in glia (*SG*, *repoGAL4>GFP*) and stained larvae with α-GFP and α-LACZ antibodies (at ZT2). To identify cortex glia, we considered the position (underneath the perineurial and subperineurial glia that surround the CNS) and the morphology of the GFP-positive cells. Fig. 4e shows that SG is detectable in cortex glia, supporting the notion that PER is expressed in these cells.

**Fig 4.**
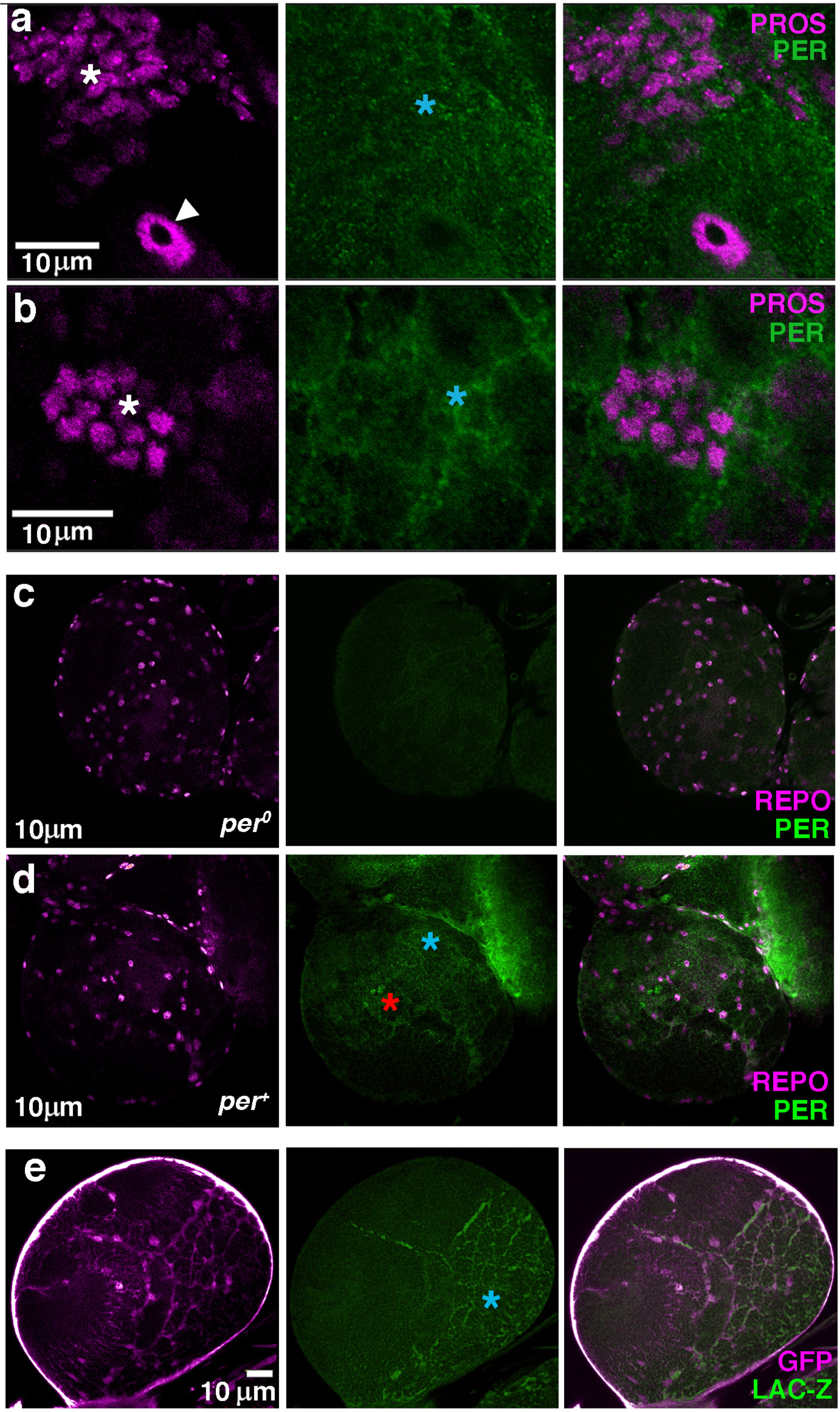
PER is expressed in glia. a, b) Optical slices (confocal) showing anti-PROS (magenta) and anti-PER (green) in brain lobes (BLs) of 3^rd^ instar larvae. The white arrowhead shows a type I neuroblast (NB I; note the asymmetric anti-PROS cytoplasmic staining) while the white asterisks indicate ganglion mother cells (GMCs; anti-PROS nuclear staining). Anti-PER does not overlap with anti-PROS but labels surrounding areas (blue asterisks). Size bars = 10 μm. ZT=23. c, d) Optical slices (confocal) showing anti-REPO (magenta) and anti-PER (green) in the BLs of 3^rd^ instar *per^0^* (c) and *per*^+^ (d) larvae. Anti-REPO stains the nucleus of glial cells. In *per*^+^, anti-PER labels the nucleus of clock neurons (red asterisk) and shows weak, diffuse staining (blue asterisk). Size bars = 10 μm. ZT=23. e) A confocal optical section showing the expression of the PER reporter SG in the BL of a 3^rd^ instar larva. SG is combined, by crossing, to *repo-GAL4>GFP* to identify glia through GFP immune reactivity. Note that the expression of SG does not cycle^37^. In the focal plane shown, anti-GFP staining (magenta) reveals cortex glia, a type of glia that envelops NBs and their progeny providing niche function. Anti-LacZ (green) shows overlapping staining (blue asterisk). Size bar = 10 μm. ZT=2. [Note: magenta and green are pseudo colors; anti-GFP and anti-LACZ were imaged in the green – Alexa488 – and red – Texas Red – channel, respectively].

### NON-AUTONOMOUS GENOTOXIC EFFECTS OF per^0^

Since we observed expression of PER and of its reporter SG in glia, we asked whether chromosome aberrations, which are detected in mitotic cells, are caused by a non-autonomous mechanism. The overexpression of PER in glia, in an otherwise *per^0^* background (*repoGAL4*>PER, *per^0^*), was sufficient to rescue chromosome aberrations (Fig. 5a). We then used a CRISPR/Cas9 approach to carry out the opposite manipulation^38^. We induced somatic mutations in the *per* gene of wild type larvae by driving the expression of CAS9 and *per* gRNA in glia (*repoGAL4*>*gRNAper*, CAS9). We observed a significant increase in the proportion of chromosome aberrations compared to the control (*repoGAL4*>*gRNAacp,* CAS9) that targets CAS9 to *acp98A*, a gene expressed exclusively in the male accessory gland^38^ (Fig. 5b).

**Fig 5.**
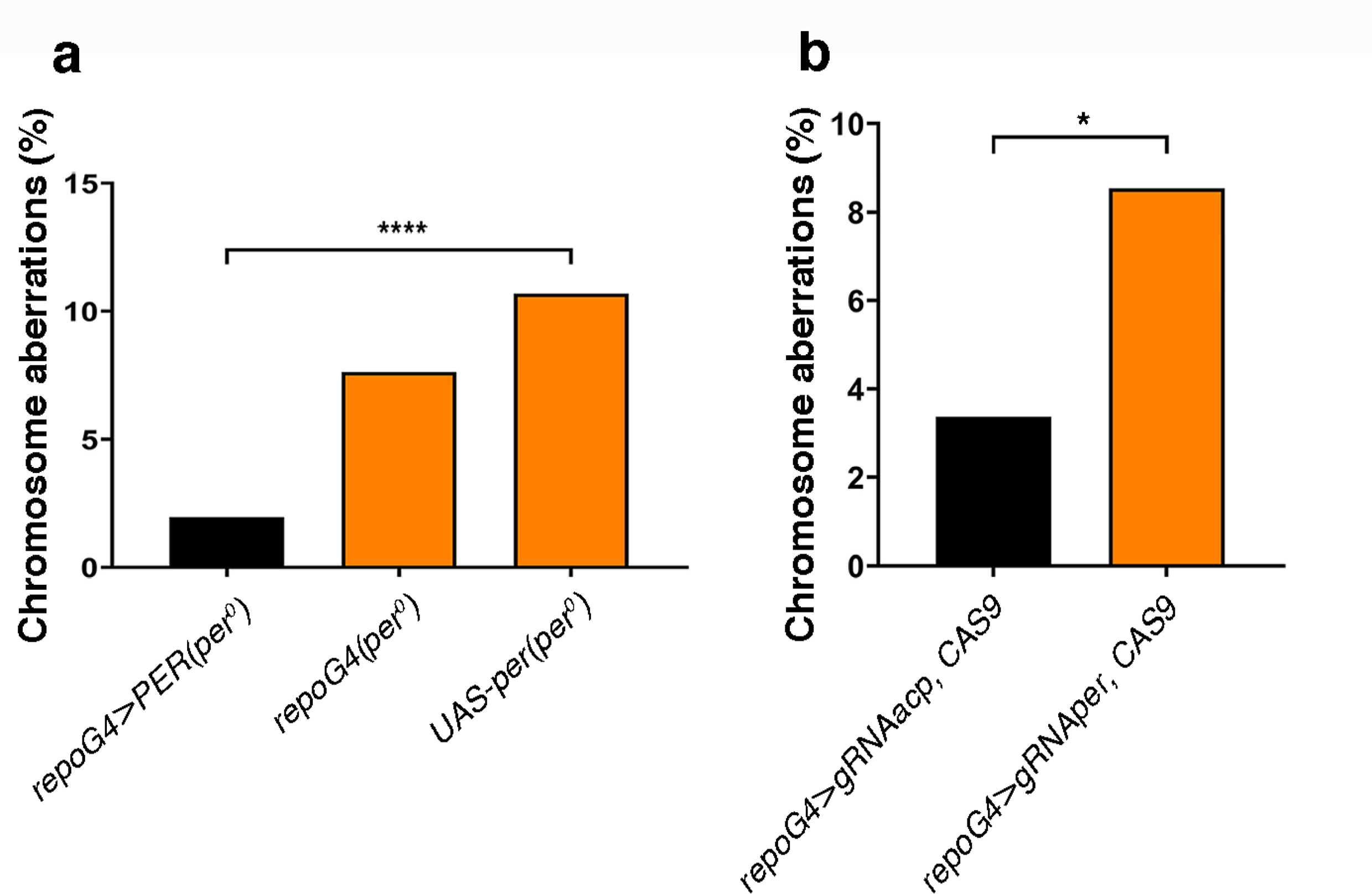
Chromosome aberrations are contingent with the lack of PER expression in glia. a) The overexpression of PER (*UASper16*) in glia [*repo-GAL4*>PER (*per^0^*)] drastically reduced the frequency of aberrations in an otherwise *per^0^* background. Chi-square=42.21, df=2, ****P<0.0001. Total number of metaphases scored (from left to right), N = 612, 302, 1021. ZT1. b) The knock-out of *per* (with CRISPR/Cas9) only in glia (*repo-GAL4*> *gRNAper*, CAS9) was sufficient to trigger chromosome aberrations. *repo-GAL4*>*gRNAacp*, CAS9 *vs*. *repo-GAL4*> *gRNAper*, CAS9, Fisher’s exact test, *P=0.0161. Total number of metaphases scored, N = 386, 164. ZT=1.

### per^0^ MAY AFFECT THE CHROMATIN LANDSCAPE OF NEURAL PROGENITOR CELLS

ROS stress can induce reprogramming of stem cells^39^. One of the possible mechanisms is DNA damage leading to H2AV phosphorylation (i.e., γ-H2AV) and then activation of the Poly(ADP-ribose) Polymerase 1 (PARP1) protein. This path causes loosening of compacted chromatin to allow DNA repair but has knock-on effects on transcription and may impact the differentiation status of the cells^40^. Additionally, in mammalian fibroblasts the knock-out of the three *Per* genes (*Per1-3*, which correspond to the single *per* in *Drosophila*) causes a reduction in the deposition of H2AZ (a homologous of H2AV) resulting in greater genome accessibility and persistent DNA damage^10^. On this premise, we decided to assess whether there may be relaxation of compacted chromatin in *per^0^*.

H3K9me3 (tri-methylated histone 3 at Lys9) is an epigenetic mark of constitutive heterochromatin and SUPRESSOR OF VARIEGATION3-9 [SU(VAR)3-9, a histone methyltransferase] and HETEROCHROMATIN PROTEIN 1 (HP1, a structural protein that binds to H3K9me3) are fundamental regulators of its formation and maintenance^41^. We used *timGAL4* to overexpress SU(VAR)3-9 or HP1 in *per^0^* larvae and we measured chromosome aberrations. Both manipulations reduced the proportion of aberrant metaphases (Fig. 6a, b). This suggests that the genotoxic effects we uncovered, may be triggered by an anomalous chromatin landscape in the cells that is caused by lack of PER or by a defect of the clock as a whole.

**Fig 6.**
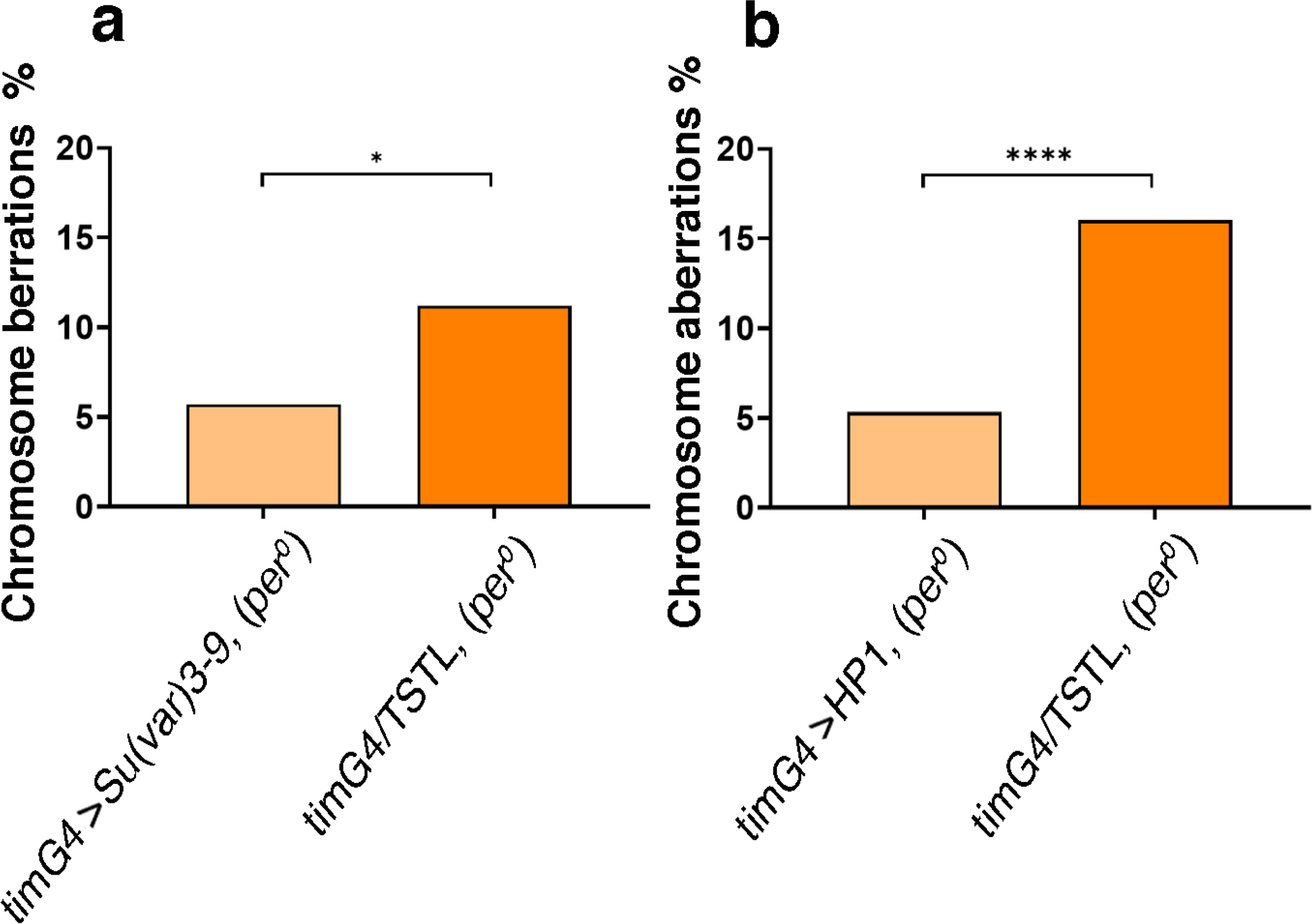
Promoting chromatin compaction reduces chromosome aberrations in *per^0^*. The overexpression of SUR3-9 (a) or HP1 (b) using the pan-circadian *tim-GAL4* driver, drastically reduced the frequency of aberrations in *per^0^*. a) *tim-GAL4*>SUR3-9 (*per^0^*) *vs*. *tim-GAL4/TSTL* (*per^0^*), Fisher’s exact test, *P<0.0129. Total number of metaphases scored, N = 388, 276. ZT=1. b) *tim-GAL4*>HP1 (*per^0^*) *vs*. *tim-GAL4/TSTL* (*per^0^*), Fisher’s exact test, ****P<0.0001. Total number of metaphases scored, N = 417, 362. ZT=1. [Note: *TSTL*, *Triplo-Sensitive & Triplo-Lethal* is a chromosomal arrangement that may be used to maintain genetic stability in crosses].

## Discussion

Previous reports had highlighted metabolic deficiencies and sensitivity to ROS in *per^0^* (adult) flies^25^. We investigated ROS levels in larvae using MitoSox Red, a fluorogenic compound targeted to the mitochondria. We detected an increased ROS burden in the CNS of *per^0^* larvae compared to the control (Fig. 1a, b). Surprisingly, using high resolution respirometry (Oroboros) we were unable to identify defects in respiration in *per^0^* larvae (Fig. S2). We analysed mitochondrial respiration in whole body extracts, which may have masked CNS-specific differences. Additionally, during the larval stages *Drosophila* relies more on glycolysis than OXPHOS for energy production^30^. Besides, larval mitochondria express Uncoupling Protein 4C (UCP4C), which leaks some of the H^+^ gradient required to generate ATP to produce heat instead. In nature, this is essential for the growth of larvae at suboptimal temperatures^31^ and may have contributed to masking differences between *per^0^*and the wild type. Thus, we looked at respiration in adult flies and found defects in OXPHOS and electron transport in *per^0^* (Fig. 1d). We suggest that mitochondria may be defective in *per^0^* larvae also, and generate high levels of ROS. This would explain the genotoxic effects we observed (Fig. 2 a-d). Indeed, by buffering ROS with vitamin C (a scavenger) or by reducing their production with AOX (an alternative oxidase that ‘shortcuts’ the electron transport chain diverting electrons from the production of ROS), we were able to reduce dsDNA damage and to restore the frequency of chromosome aberrations to wild type levels (Fig. 3a-g). Notably, we expressed AOX using *timGAL4*. This suggests that the cells important for phenotypic rescue express both *per* and *tim*, which qualifies them as putative clock cells.

In *Drosophila* the role that the circadian clock, or its components, have on development, is little explored and poorly understood. In embryos, PER and TIM are expressed in many cells of the CNS from embryonic stage 12 (ES12), but only at the end of ES16 they seem to overlap in about 20 cells of the protocerebrum^13, 14^. In larvae, the pattern of expression is even more uncertain. There is agreement that in each brain lobe (BL) there are 9 *bona fide* clock neurons expressing rhythmic PER and TIM^42-45^. However, whether and when the other clock neurons (which are more than 75 per hemisphere in the adult) become differentiated in larvae and whether they express PER and/or TIM, is not clear. For instance, while confocal microscopy showed that some dorsal clock neurons become evident in late 3^rd^ instar larvae but having little PER expression and no cycling^45^, using a spatially-restricted PER-LUCIFERASE reporter suggested that PER is rhythmically expressed in these neurons from the end of embryonic development^46^. This and other discrepancies may be explained by the low level of expression of the native proteins, by the complex expression pattern of the reporters, and by the lack of independent markers to identify clock cells^42-45^.

In this work, we have shown α-PER immune labelling and expression of a ‘classic’ PER reporter, SG, in larval glia cells. Kaneko and Hall (2000, see Fig.6 therein) had identified PER expression in larval glia using a different reporter (*perGAL4*>TAU), which provides independent confirmation for our finding^43^. Here, by combining SG with GFP expression in glia (*SG*, *repoGAL4*>GFP) and by comparing the position and the morphology of the GFP-positive cells, we were able to identify some putative PER-expressing glia cells as cortex (Fig. 4d). These are large, web-like cells, born during the mid-embryonic stage, that surround and support the NBs and their lineages functioning as ‘niche’^36^. Liu et al. (2015) did not detect PER staining in glia^45^. However, a further validation of our observations comes from the fact that manipulating glia either by overexpressing PER (*repoGAL4*>PER, *per^0^*) in *per^0^* or by mutating *per* (*repoGAL4*>*gRNAper*, CAS9) in a *per*^+^ background, we were able to rescue or to induce, respectively, chromosome aberrations observed in *per^0^* larvae (Fig.5 a, b).

In *Drosophila*, H2AV is homologous to both histone variants H2AX and H2AZ found in mammals. As for H2AX, phosphorylation of H2AV at Ser 137 (Ser 139 in H2AX) signals the presence of dsDNA breaks and leads to the recruitment of repair proteins at the break sites. Moreover, like H2AZ, H2AV regulates transcription and chromatin structure^33^. Indeed, both H2AV and H2AZ are found at the promoter region of active genes supporting their transcription, but also in correspondence of facultative and constitutive heterochromatin contributing to their establishment and repressive function^33, 47, 48^. Besides, both H2AV and H2AZ interact with insulator proteins and regulate their deposition across the genome^49, 50^. Insulators mediate the 3D-organisation of chromatin by promoting the formation of topological associating domains (TADs) that establish ‘locally regulated areas’ of gene expression^51, 52^. The phosphorylation of H2AV to γ-H2AV leads to the activation of PARP1, an enzyme that transfers ADP-ribose units from NAD^+^ to target proteins. This promotes the opening of chromatin by destabilizing pre-existing protein complexes, which facilitates transcription and genotoxic stress responses^40^. Finally, in mammalian fibroblasts a triple *Per1-3* knock-out (that removes the three homologous *Per* genes) causes reduced deposition of H2AZ, greater genome accessibility and persistent DNA damage^10^. Overall, these findings suggest that we may find epigenetic dysregulation in *per^0^*. Thus, we tested whether *per^0^* larvae may incur a reduction in the compaction of the chromatin by overexpressing proteins that could counteract such a phenomenon. We used *timGAL4* to overexpress SU(VAR)3-9 and HP1 in all putative clock cells in *per^0^* larvae. SU(VAR)3-9 and HP1 are involved in the formation and maintenance of H3K9me3, a fundamental component of constitutive heterochromatin^41^. Both manipulations rescued the chromosome aberration that were otherwise observed in *per^0^*mutants (Fig.6 a, b).

In summary, we have uncovered novel phenotypes in third instar larvae of *per^0^* mutants, namely increased dsDNA breaks in the CNS and a high frequency of chromosome aberrations in mitotic precursors of neuronal cells. For the latter, we have shown that PER expression in glia is necessary and sufficient to avoid such an effect. We have shown that ROS are a main trigger for these phenotypes and that *per^0^* is at the origin of mitochondrial dysfunctions that generate ROS. Finally, we suggest that *per^0^* larvae may experience widespread decompaction of the chromatin, resulting in abnormal epigenetic regulation. Our work shows how a fundamental constituent of the circadian clock, perhaps the clock itself, can influence metabolism and chromatin structure to regulate development, which perhaps points to a fundamental function of the clock that goes beyond rhythmicity.

## Materials and methods

### *Drosophila* strains and maintenance

Flies were maintained at 25°C in a 12h: 12h light-dark (LD) cycle on maize/glucose/yeast/agar (6.3/6.9/4.4/0.5 %) medium using propionic acid as a mold inhibitor.

We used the following stocks, *Canton-S* (*CS*), *per*^0^ (in *CS* background), yw; *per*GAL4, and *SG^10^*; *tim^0^* (refs: 42, 43, 37; from Jeff Hall, Brandeis University, USA); *per*^0^;; UAS-*per*^16^ (ref: 53; from François Rouyer, CNRS, France); *yw; tim*GAL4 (ref: 54; from Patrick Emery, UMass, USA); *w^1118^;; repoGAL4/TM3* (#7415, Bloomington *Drosophila* Stock Center, Indiana, USA); *w^cs^; UASgRNAacp/CyO; UASCAS9/TM3* and *w*^cs^; *UASgRNAper/CyO*; *UASCAS9/TM3* (ref: 38; from Mimi Shirasu-Hiza, Columbia University, USA); *w^1118^; dpnEEGal4/CyO; TM3/TM6* (ref: 55; from Tzumin Lee, Janelia Research Campus, USA); *w^1118^; UAS-AOX* (ref: 35; from Howard T. Jacobs, Institute of Medical Technology and Tampere University Hospital, Finland); *UASSu(var)3-9GFP*/*T(2;3)TSTL* and *w^1118^; UAS-Hp1/CyO* (from Lucia Piacentini, ‘La Sapienza’ University, Italy).

### Antioxidant feeding

Pure vitamin C was diluted in water (H_2_O) and added to the standard feeding medium to a final concentration of 40mM^34^. In vehicle only controls the same amount of water was added to the medium. Adult flies were transferred to fresh tubes every 2-3 days.

### Immuno-staining and confocal microscopy

For whole mount immunolabelling, the CNS of 3^rd^ instar larvae were dissected in cold phosphate buffer saline (PBS) and fixed in 4% paraformaldehyde (PFA) in PBS for 30 minutes at room temperature (RT). Samples were washed three times in PBS with 0.5% Triton X 100 (0.5% PBS-Tx) for 15 min at RT. Then, they were blocked with 10% normal goat serum in 0.5% PBS-Tx for 1 hour at RT and immunolabelled with primary antibodies (diluted in fresh blocking solution) at 4°C overnight. Samples were washed three times in 0.5% PBS-Tx for 15 min at RT and incubated with secondary antibodies (diluted in 0.5% PBS-Tx) for 3.5h at RT in the dark. Samples were mounted on slides with antifade medium (3% propyl gallate, 80% glycerol, 20% 1xPBS, pH 8.5). Five to ten brains were scored for each experiment. Observations were performed either on an Olympus FV1000 or on Zeiss LSM 780 confocal microscope. Microscope, lasers, filters, and all other settings remained constant within each independent experiment. Images were processed using FIJI (release 1.54f)^56^.

### Antibodies

Primary: chicken α-GFP (1:1000, Abcam #ab13970), mouse monoclonal α-LacZ (1:1000, Promega #Z3781), mouse monoclonal α-γH2Av (1:10, DSHB #UNC93-5.2.1), mouse monoclonal anti-Repo (1:15, DSHB #8D12), mouse monoclonal anti-Prospero (1:10, DHSB #MR1A), rabbit α-PER c-300, (1:50; Santa Cruz Biotech).

Secondary: goat anti-mouse AlexaFluor568 (1:1000, Invitrogen), goat anti-mouse Texas Red (1:400, Jackson ImmunoResearch), goat anti-rabbit Cy2 (1:400, Jackson ImmunoResearch) goat anti-chicken AlexaFluor488 (1:400, Invitrogen).

### Measurement of mitochondrial ROS

Central nervous systems (CNSs, each consisting of two brain lobes – BLs – and one ventral nerve cord – VNC –) from male 3^rd^ instar larvae were dissected in cold phosphate buffer saline (PBS) and incubated with 5 μM of MitoSOX Red (Invitrogen) for 30 min at room temperature (RT). After incubation, CNSs were washed 3x5 min with PBS at RT. Then, they were fixed with 4% PFA for 20 min at RT, washed 3x5 min with PBS and mounted in antifade (3% propyl gallate, 80% glycerol, 20% 1xPBS, pH 8.5). Samples were imaged immediately with an Olympus FV1000 confocal laser scanning microscope. Images were acquired as z-stacks through the entire thickness of the BLs and (separately) the VNC using a 20× UPlanSApo Olympus objective. Total fluorescent intensities for the BLs (the two were averaged for each individual) and the VNC were measured using Fiji (release 1.54f)^56^.

### Mitotic chromosome preparations

Mitotic chromosomes from the CNS were prepared as previously published^57^. Briefly, CNSs were dissected from male 3^rd^ instar larvae in physiological solution (NaCl 0.7%), transferred to hypotonic solution (sodium citrate 0.5%) for 8 min and then moved to a drop of fixing solution (methanol: acetic acid: water = 5.5: 5.5: 1) for 30 seconds. Five CNSs were transferred to five drops of 45% acetic acid on a siliconized coverslip. A non-siliconized slide was lowered on the coverslip, the ‘sandwich’ was inverted and squashed between two sheets of blotting paper for 1 min. Slides were frozen in liquid nitrogen and the coverslips were ‘flung off’ with a razor blade. The slides were immersed in 100% ethanol for 5 min. Then, they were washed in 1XPBS for 10 min and counterstained with DAPI solution (0.05 μg/ml of 4ʹ, 6-diamidine-2-phenylindole dihydrochloride in 2XSSC) for 5 min. Samples were mounted in antifade medium (2.3% DABCO – 1, 4-diazabicyclo[2,2,2]octane –, 20mM Tris-HCl pH 8, 90% glycerol) and imaged on a Nikon fluorescent microscope equipped with a cooled CoolSnap CCD camera (Photometrics). Images were processed using Adobe Photoshop.

### Immunofluorescence on CNS-squash preparations

We followed a previously published protocol^57^. Briefly, CNSs were dissected from male third instar larvae in physiological solution (NaCl 0.7%), transferred to hypotonic solution (sodium citrate 0.5%) for 8 min, and then fixed in 45% acid acetic, 2% formaldehyde for 10 min, before being squashed for 1 min in the same solution. Slides were frozen in liquid nitrogen and the coverslips were removed. Slides were transferred to 1XPBS for 5 min, permeabilized in 1% PBS-Tx for 10 min, and blocked in 1X PBS, 1% BSA, for 30 min before incubation with mouse α-γ-H2AV antibodies (DSHB #UNC93-5.2.1) diluted 1:5 in 1X PBS, 1% BSA. Incubation with the primary antibodies was carried at room temperature for 1 h and then at 4 °C overnight. Samples were washed 3×5 min in 1X PBS. The secondary antibodies, goat anti-mouse Cy3 (Jackson ImmunoResearch), were diluted 1:400 in 1X PBS, 1% BSA and incubated at room temperature for 2 h. Samples were washed 3×5 min in 1X PBS, counterstained with DAPI solution (0.05 μg/ml of 4ʹ, 6-diamidine-2-phenylindole dihydrochloride in 2XSSC) for 5 min and then mounted in Vectashield H-1000 (Vector Laboratories). Samples were imaged using a fluorescent Nikon microscope equipped with a cooled CoolSnap CCD camera (Photometrics). The exposure time was kept constant across all samples. The two fluorescent signals (from DAPI and Cy3), were recorded separately as gray scale digital images. Images were pseudo coloured and merged using Adobe Photoshop.

### Mitochondrial oxygen consumption measurements

High resolution respirometry measures were determined with an Oroboros Oxygraph-2k (Oroboros Instruments, Austria). Mitochondrial leak (L), oxidative phosphorylation (OXPHOS) and electron transport system (ETS) capacities were quantified using a previously described substrate-uncoupler-inhibitor-titration (SUIT) protocol^58^, with the following modifications. Only males, *per*^+^ and *per^0^*, were used for the analyses. The two genotypes were obtained, respectively, by crossing female *CS* to males *per^0^* and *vice versa*. For each sample, ten 3-5 day old flies (or ten 3^rd^ instar larvae) from the same genotype, were homogenized together in 800 μl of respiration buffer MiR05 (0.5 mM EGTA, 3 mM MgCl2, 60 mM K-lactobionate, 20 mM taurine, 10 mM KH2PO4, 2 0 mM HEPES, 110 mM sucrose, and 1 g/l BSA, pH 7.1). When testing, the two chambers of the oxygraph, each containing 2 ml of MiR05, were loaded with 80 μl of homogenate, one from a *per*^+^, the other from a *per^0^* sample. The chambers were extensively washed between tests, inverting the loading at each cycle.

### Statistics and reproducibility

Data have been plotted and analysed using Graphpad Prism 9.5.1. We employed Chi-squared, Fisher’s exact test, Shapiro-Wilk test, Kolmogorov-Smirnov test, Mann-Whitney test, and Two-Way ANOVA with Tukey’s post-hoc analyses, as appropriate. All statistical tests used were two-tailed. Sample sizes are indicated in figures and/or legends.

## Acknowledgements

We acknowledge the following funding from *Sapienza* University*, Dottorato in Genetica e Biologia Molecolare* (NCR & LL), *Progetti di Ateneo* (LF), Funding for International Research Placements (DR) and Visiting Professorship (ER). ER thanks Cinchron (Marie Skłodowska-Curie Actions grant agreement 765937) for support. We acknowledge the Developmental Studies Hybridoma Bank, created by the NICHD of the NIH and maintained at The University of Iowa, Department of Biology, Iowa City, IA 52242 for the monoclonal antibodies anti-Repo (developed by C. Goodman), Prospero (developed by C. Q. Doe), and γH2Av (developed by R. S. Hawley).

## Contributions

Conceptualization, NCR, ER and LF; Methodology, NCR, MM, ER and LF; Investigation and analyses, NCR, MM, AM, LL, DR, ER, LF; Writing – original draft, NCR, ER and LF; Writing - Review and editing, NCR, MM, AM, LL, DR, ER, LF; Supervision, MM, ER, and LF; Funding Acquisition, ER and LF.

## Data availability

All data generated or analysed during this study are included in this article. This work did not generate novel materials or tools. Data and materials used are available from the corresponding authors upon request.

---

**Fig S1.**
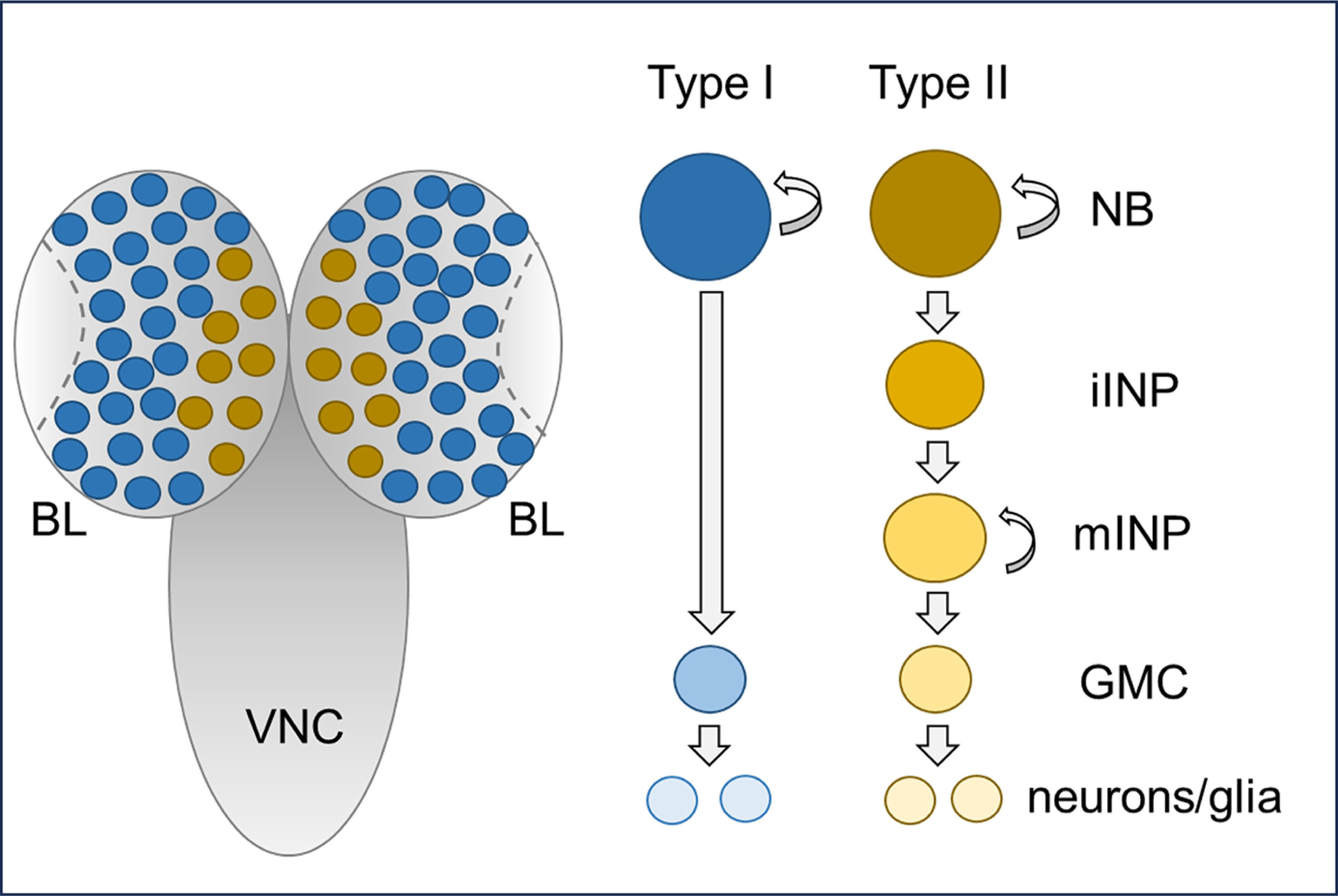
[linked to Fig1]. Schematic representation of the central nervous system (CNS) of *Drosophila* larvae. Distribution of type I (blue) and type II (gold) neuroblasts in the brain lobes. In the ventral cord, only type I neuroblasts are present (not shown). BL: brain lobe; VNC: ventral nerve cord; NB: neuroblast; iINP: immature intermediate progenitor; mINP: mature intermediate progenitor; GMC: ganglion mother cell.

**Fig S2.**
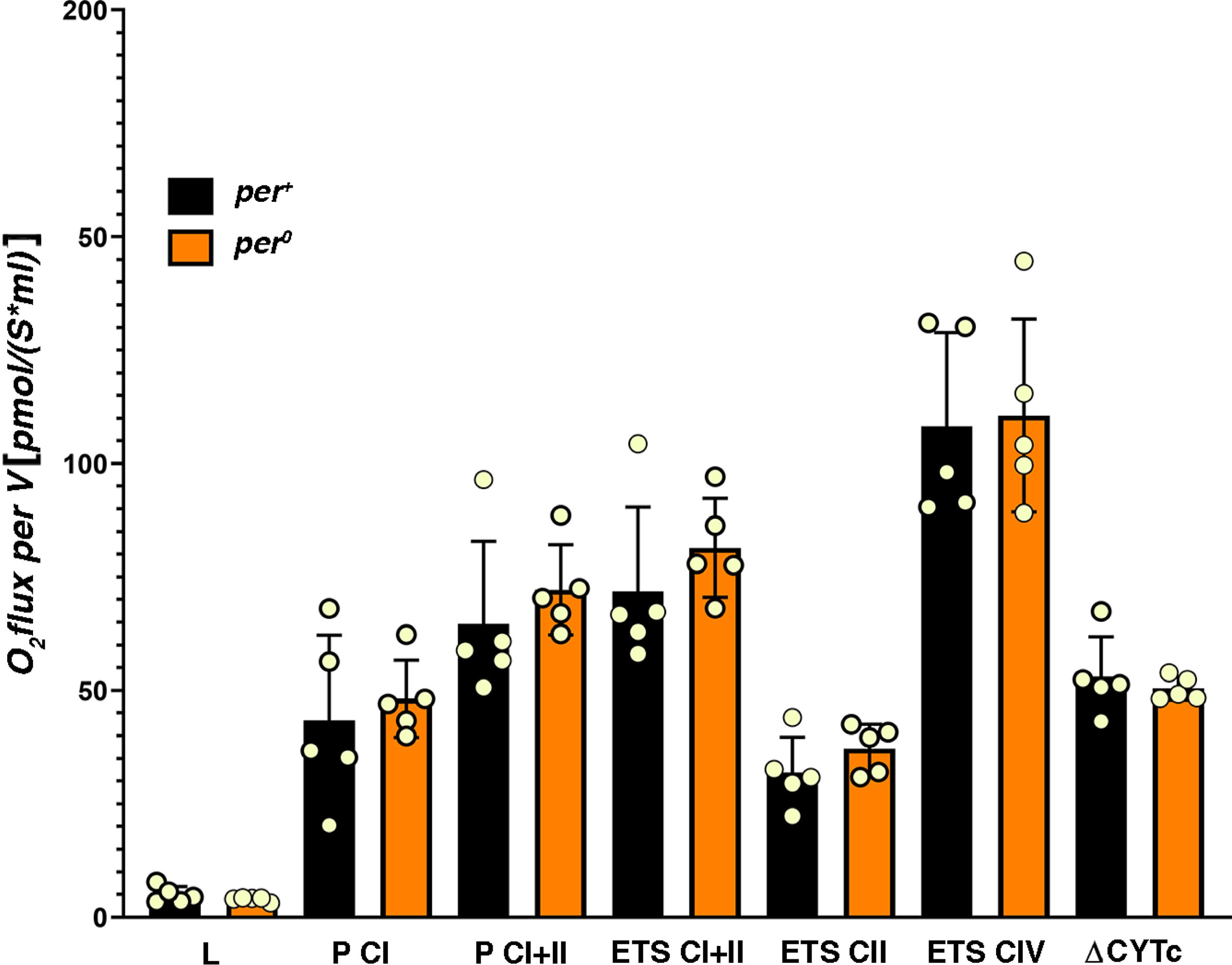
[linked to Fig1]. High resolution respirometry does not identify differences between *per*^+^ and *per^0^* in 3^rd^ instar larvae. L, mitochondrial leak (Mann-Whitney, P=0.421). P CI, OXPHOS capacity of complex I (Mann-Whitney, P=0.548). P CI + II, OXPHOS capacity of complex I plus II (Mann-Whitney, P=0.151). ETS CI + II, electron transport capacity through complex I plus II (Mann-Whitney, P=0.151). ETS CII, electron transport capacity through complex II (Mann-Whitney, P=0.310). ETS CIV, electron transport capacity through complex IV (Mann-Whitney, P=0.841). ΔCYTc, increase in ETS CIV when excess cytochrome c is added to the respiration buffer (measure of mitochondria integrity, Mann-Whitney, P=0.691). Points show individual samples. Error bars = SD. ZT=1. All samples were males obtained by reciprocal crossing (♀ *CS* x ♂ *per*^0^ and *vice versa*).

